# Frequent emergence of resistance mutations following complex intra-host genomic dynamics in SARS-CoV-2 patients receiving Sotrovimab

**DOI:** 10.1101/2023.03.01.530733

**Authors:** Rosalía Palomino-Cabrera, Francisco Tejerina, Andrea Molero-Salinas, María Ferris, Cristina Veintimilla, Pilar Catalán, Gabriela Rodríguez Macias, Roberto Alonso, Patricia Muñoz, Darío García de Viedma, Laura Pérez-Lago, the Gregorio Marañón Microbiology-ID COVID 19 Study Group

**Author notes:** **Corresponding authors:** Darío García de Viedma and Laura Pérez-Lago, Postal address: C/Dr. Esquerdo 46 (28007), Madrid, Spain, Mail. both authors contributed equally as senior authors (Darío García de Viedma, Laura Pérez-Lago).

## Abstract

The emergence of the Omicron variant of SARS-CoV-2 represented a challenge to the treatment of COVID-19 with monoclonal antibodies. Only Sotrovimab maintained partial activity, allowing it to be used in high-risk patients infected with the Omicron variant. However, the reports of resistance mutations to Sotrovimab call for efforts to better understand the intra-patient emergence of this resistance. A retrospective genomic analysis was conducted on respiratory samples from immunocompromised patients infected with SARS-CoV-2 who received Sotrovimab at our hospital between December 2021 and August 2022. The study involved 95 sequential specimens from 22 patients (1-12 samples/patient; 3-107 days post-infusion (Ct ≤ 32)). Resistance mutations (in P337, E340, K356, and R346) were detected in 68% of cases; the shortest time to detection of a resistance mutation was 5 days after Sotrovimab infusion. The dynamics of resistance acquisition were highly complex, with up to 11 distinct amino acid changes in specimens from the same patient. In two patients, the mutation distribution was compartmentalized in respiratory samples from different sources. This is the first study to examine the acquisition of resistance to Sotrovimab in the BA.5 lineage, enabling us to determine the lack of genomic or clinical differences between Sotrovimab resistance in BA.5 relative to BA.1/2. Across all Omicron lineages, the acquisition of resistance delayed SARS-CoV-2 clearance (40.67 vs 19.5 days). Close, real-time genomic surveillance of patients receiving Sotrovimab should be mandatory to facilitate early therapeutic interventions.

---

The emergence of the Omicron variant of SARS-CoV-2 variant had a major impact on the treatment of COVID-19, as this lineage was resistant to most monoclonal antibodies (mAb). Despite this, Sotrovimab, which targets an epitope in the receptor-binding domain (RBD) of the spike protein, a highly-conserved region in sarbecoviruses (1, 2), maintains partial in vitro activity against different Omicron sublineages and has been used until recently in high-risk patients infected with the Omicron variant of SARS-CoV-2.

In Spain, the administration of Sotrovimab in the context of early treatment for high-risk patients, or as compassionate use in hospitalised immunocompromised seronegative patients with severe COVID-19, was approved in December 2021 (3). This prompted an evaluation of the possible acquisition of resistance in patients who had been receiving it at our institution since its approval.

A retrospective study was conducted in 48 COVID-19 patients who received Sotrovimab (500 mg) between December 2021 and August 2022, supported by a longitudinal genomic analysis of their sequential positive samples. Twenty-eight patients (all immunocompromised) had at least one positive sample available (with sufficient viral load to be sequenced Ct ≤ 32) four days after Sotrovimab infusion, and were included in our analysis. Diagnostics and Ct determination were performed on respiratory specimens by RT-PCR (TaqPath COVID-19 CE-IVD RT-PCR kit; ThermoFisher Scientific, MA, USA). Whole genome sequencing was carried out as described elsewhere (4). Sequence analysis was performed using an in-house bioinformatics pipeline (https://github.com/MG-IiSGM/covid_multianalysis), giving sequencing data with quality scores above the threshold for 95 specimens from 22 patients (1-12 specimens per patient, 3-107 days post-infusion). The requirements for SNP calls were: frequency ≥ 5% and total depth ≥ 10X; a SNP was considered fixed when its frequency was >80% and was considered non-fixed when present at frequencies between 5-80%. Minority SNP calls (< 5%) always required visual inspection by IGV. When necessary, short tandem repeat (STR) analysis was performed to ensure that the specimens belonged to the same individual (4).

The time interval between COVID diagnosis and Sotrovimab infusion was 1-4 days in 8 cases where it was administered as early treatment due to the risk of developing severe COVID-19, 1-51 days in 13 hospitalized patients with severe COVID, and 68 days in one patient with persistent infection.

## Emergence of resistance mutations

Resistance mutations were detected in 15 patients (68%), which places our findings above the highest frequencies (55%-60%) reported to date (5, 6). The mutations identified (frequency >5%) were distributed among the four codons previously described to encode sotrovimab resistance (P337, E340, R346 and K356) and included 14 different substitutions (P337S/R/T/L/A/H, E340Q/A/D/K/V/G, R346T and K356T). Mutations in these spike protein residues have been shown to be associated with reduced neutralization by this mAb (5–8). The wide diversity of substitutions and their distribution among four codons is in agreement with other studies (5, 6), while two studies from the Netherlands (7, 9) found more homogeneous behaviour (mutations only in P337 and E340 and low diversity of substitutions).

Resistance mutations (frequency > 5%) were identified for the first time between 5-18 days post-treatment (Supplementary Table). We explored the presence of minority mutations (<5%) in nine cases with positive samples prior to the first mutation detected. We identified mutations at lower frequencies (1.6%, data not shown) in only one case (patient 15), preceding in eight days the initial detection of mutations in this case. On the other hand, the longest period until the first detection of a resistance mutation, among the cases without mutations in the 1-5 day period, was 16 days (patient 4). Other studies have reported later onset of resistance, in the range of 14-21 days (5, 8), 6-13 days (10) and up to day 31 (9) post-treatment. However, the robustness of some of these dates is limited by the availability of sequenced specimens prior to the first one with a mutation.

## Dynamics of resistance mutations

Given that Sotrovimab has a median terminal half-life of 56.5 days post-infusion (2), a prolonged analysis of the evolution of mutations is required as soon as the first one has been identified. Our study is the most exhaustive within-patient sampling effort, covering an observation window of up to 107 days post-infusion and including up to 12 sequential positive specimens per patient.

In 12 cases, at least one of the mutations observed was eventually fixed (Figure). All 14 mutations that became fixed corresponded to either P337 or E340 (5 and 9, respectively), with the most frequent being E340K (4 times), followed by E340D and P337S (3 times each), E340Q (twice) and P337L and P337R, identified only once each. These substitutions are also found at high rates in other studies (5–8, 10). More than one different fixed substitution never co-occurred in the same specimen. However, in two cases (patients 1 and 10), two different fixed substitutions at the same amino acid were observed at two different time points (E340D/K, and P337S/R, respectively, Figure), a phenomenon previously described by other authors (6, 8).

**Figure.**
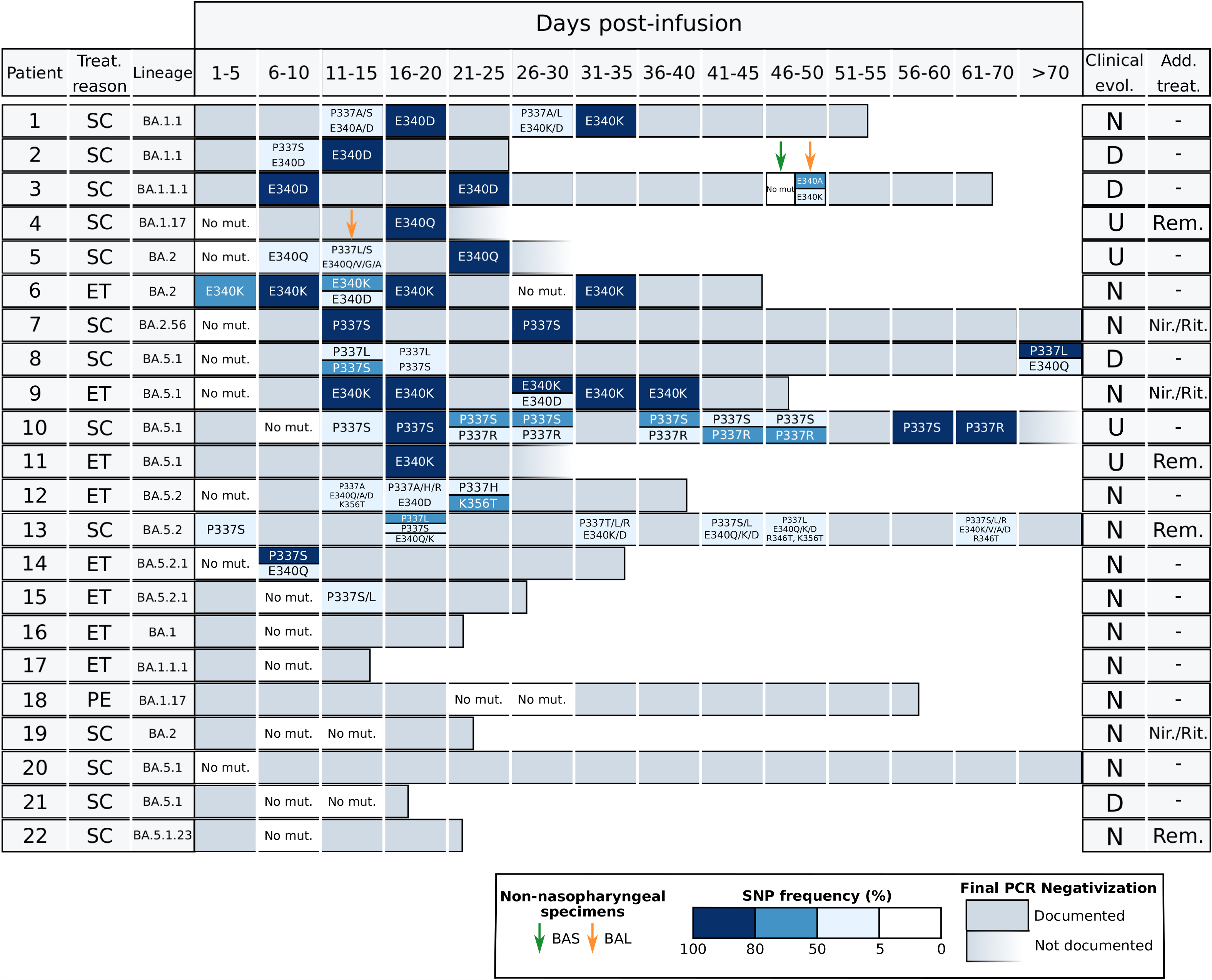
Sotrovimab resistance mutations identified in the patients in study. Each horizontal bar corresponds to the RT-PCR positivity period for each patient. For the specimens which were sequenced we indicated if we detected or not mutations and the colour gradient indicates the allele frequencies for the SNPs identified. The reasons for Sotrovimab treatment: SC (severe COVID), ET (early treatment) or PE (SARS-CoV-2 persistence) and the clinical evolution: D (died), N (negativization documented by RT-PCR), U (undefined negativization due to lack of a negative PCR) are indicated. Those patients with additional anti-viral treatment (Add. Treat) other than Sotrovimab are specified: Rem (remdesivir), Nir./Rit. (nirmatrelvir/ritonavir, paxlovid). BAL/BAS specimens are highlighted by arrows.

Regarding non-fixed substitutions (frequencies 5-80%), these were found in some of the specimens in 8 of the 12 cases in which some mutations eventually became fixed, and in another three cases in which only non-fixed mutations were detected throughout the analysis. The diversity of non-fixed substitutions was remarkably high and involved all four codons where mutations were identified (P337A/H/S/L/T/R, E340Q/A/V/D/K, K356T, R346T). The homogenous composition observed in most of the specimens with fixed mutations contrasted with the greater within-sample diversity often identified in non-fixed substitutions (either in the same codon or even in more than one codon, Figure). This diversity was highest in two of the patients (patients 12 and 13) with non-fixed mutations during the positivity period; 7 and 11 different amino acid changes, respectively, were identified (Figure). For both fixed and non-fixed mutations, we observed either a transient presence (in only one specimen) or a more prolonged presence over several sequential specimens (Figure). The data taken together illustrate the highly dynamic clonal behaviour of SARS-CoV-2 under Sotrovimab pressure.

Our study is the first to include strains of the BA.5 lineage in the genomic analysis of resistance to Sotrovimab, with the majority of cases under study (50%) belonging to this lineage (7 to BA.5.1* and 4 to BA.5.2*); the remaining variants involved were BA.1 and BA.2 (7 and 4 cases, respectively; Figure). The lack of differences in the dynamics of sotrovimab resistance acquisition found between BA.1 and BA.2 in previous studies can now be extended, based on our initial observations, to BA.5 variants. The Delta lineage has also been found to share similar patterns of acquisition of Sotrovimab resistance mutations (10). Our data support the inclusion of cases infected with the BA.5 lineage among those that could face therapeutical challenges when receiving sotrovimab, as the genomic findings for BA.1/BA.2 are equivalent to those for BA.5 based on general patterns such as the percentage of cases that acquired resistance (73% in BA.5 cases and 64% non-BA.5 cases) and other qualitative findings, such as the fixation of mutations, the coexistence of several different substitutions, the complexity of the clonal dynamics or the rapidity of detection of the emergence of resistance (5 days for both BA.5 and BA.1/BA.2).

## Resistance in non-nasopharyngeal specimens

In addition to the predominantly nasopharyngeal specimens included in our study, we had the opportunity to analyse lower respiratory specimens (bronchoalveolar lavage (BAL) or bronchial aspirate (BAS)) from two of the patients with resistance detected in NP specimens (patients 5 and 3). In patient 5, E340Q was the only mutation identified in two sequential NP specimens, whereas a greater diversity of mutations was observed to coexist in a single BAL sample (two different P337 substitutions and another four in E340; Figure). In patient 3, with E340D fixed in two sequential NP specimens, another two different non-fixed substitutions in the same codon (E340K (29%) and E340A (67%) were observed in the BAL specimen (Figure). Unexpectedly, no mutations were identified in the BAS specimen. The compartmentalized microevolution of microorganisms in different respiratory samples has also been described for SARS-CoV-2 (11), but this is the first description in Sotrovimab resistance.

## Microbiological and clinical evolutionary aspects

When the effect of the acquisition of resistance on the evolution and outcome of patients was analysed, a significant increase in the mean number of days to viral clearance was observed in patients in whom resistance mutations had been identified (40.67 days) versus those without resistance mutations (19.5 days; p=0.003 in the analysis of variance on the linear model after removing outliers detected by the Grubbs test), as reported in other studies (7, 9).

Three of the patients who acquired resistance mutations (patients 2, 3, and 8; all three received Sotrovimab due to severe COVID-19) died from COVID. Time from treatment to death was 25, 65 and 114, respectively. In all three, specimens with a diversity of substitutions and others with a fixed mutation were identified (P337L in one case and E340D in the remaining two) (Figure). Patients 2 and 8 died twelve and seven days, respectively, after their last positive RT-PCR test and both had high viral loads (around Ct 16), suggesting that they probably did not clear the virus. Patient 3 died 43 days after the last NP positive RT-PCR test (Ct 26) and no subsequent RT-PCRs were performed, so that viral clearance before death was not evaluated.

The BA.5 lineage was not associated with a differential pattern in either patient outcome (18% of patients died in both groups) or mean time to viral clearance (for BA.1/2 lineages: 49 days for patients with resistance vs 19 days for patients without resistance; for BA.5 lineages: 36.5 days and 21 days for patients with and without resistance, respectively).

Seven patients received additional treatments for COVID-19 other than Sotrovimab; three received nirmatrelvir/ritonavir and four received remdesivir (in both treatments, two patients received antivirals as early treatment, and viral persistence was maintained). None of these patients died from COVID and no decrease in time to viral clearance (20 to 124 days) was found. In four of the five cases where mutations were identified, one evolved to fixation; in two of the remaining cases, no mutations were detected. The limited number of patients in the study receiving treatment with two antiviral drugs limits the statistical power of our observations. In addition, the majority of these patients received treatment as sequential therapy, with Sotrovimab being used as salvage therapy after microbiological failure of early treatment with nirmatrelvir/ritonavir or remdesivir. Thus, the suggested possible benefits of combination treatments of small-molecule antiviral drugs associated with Sotrovimab in other studies cannot be evaluated in this cohort of patients (8).

## Conclusions

Our study found the highest rates of acquisition of resistance mutations reported to date in immunocompromised patients receiving Sotrovimab. The increased diversity of lineages in our analysis includes for the first time Omicron BA.5, as well as BA.1 and BA.2, among the variants that make treatment with Sotrovimab challenging. Our genomic analysis on a high number of sequential specimens offers a detailed snapshot of the highly dynamic and complex mutational pathways followed by SARS-CoV-2 under the selective pressure of Sotrovimab, even leading to an asymmetrical compartmentalized distribution of mutations at different sites. Acquisition of resistance leads to a delay in the time needed to clear the virus, with all the potential clinical implications that this implies. Close real-time genomic surveillance of patients receiving Sotrovimab should be mandatory to allow for timely therapeutical interventions as soon as resistance is detected.

## Acknowledgements

This work was supported by the Instituto de Salud Carlos III (PI21/01823) together with the FEDER fund “A way of making Europe”, and by the ECDC (2021/PHF/23776), and IiSGM (2021-II-PI-01). Miguel Servet contract (CPII20/0000175) to LPL and PFIS contract to (FI20/00129) CRG. We would like to thank the Genomics Unit of our institution for all the support in the sequencing runs and Janet Dawson for her help in editing and proofreading.

## Ethical statement

The study was approved by the research ethics committee of Gregorio Marañón Hospital (REF: MICRO.HGUGM.2020-042). Due to the retrospective nature of the study, informed content was not required by the Ethic Committee from our institution.

### Gregorio Marañón Microbiology-ID COVID 19 Study Group

Alcalá (Luis), Aldámiz (Teresa), Alonso (Roberto), Álvarez-Uría (Ana), Bermúdez (Elena), Bouza (Emilio), Buenestado-Serrano (Sergio), Burillo (Almudena), Carrillo (Raquel), Catalán (Pilar), Cercenado (Emilia), Cobos (Alejandro), Díez (Cristina), Escribano (Pilar), Estévez (Agustín), Fanciulli (Chiara), Galar (Alicia), García (Mª Dolores), García de Viedma (Darío), Gijón (Paloma), Guillén (Helmuth), Guinea (Jesús), Herranz (Marta), Irigoyen (Álvaro), Kestler (Martha), López (Juan Carlos), Machado (Marina), Marín (Mercedes), Martín-Rabadán (Pablo), Molero-Salinas (Andrea), Montilla (Pedro), Muñoz (Patricia), Padilla (Belén), Palomino-Cabrera (Rosalía), Palomo (María), Pérez-Granda (María Jesús), Peñas-Utrilla (Daniel), Pérez-Lago (Laura), Pérez (Leire), Reigadas (Elena), Rincón (Cristina), Rodríguez (Belén), Rodríguez (Sara), Rodríguez-Grande (Cristina), Rojas (Adriana), Ruiz-Serrano (María Jesús), Sánchez (Carlos), Sánchez (Mar), Sanz-Pérez (Amadeo), Serrano (Julia), Tejerina (Francisco), Valerio (Maricela), Veintimilla (Mª Cristina), Vesperinas (Lara), Vicente (Teresa), de la Villa (Sofía).

